# Songbird organotypic culture as an *in vitro* model for interrogating sparse sequencing networks

**DOI:** 10.1101/164228

**Authors:** Jun Shen, Todd A. Blute, William A. Liberti, William Yen, Derek C. Liberti, Darrell N. Kotten, Alberto Cruz-Martín, Timothy J. Gardner

## Abstract

Sparse sequences of neuronal activity are fundamental features of neural circuit computation; however, the underlying homeostatic mechanisms remain poorly understood. To approach these questions, we have developed a method for cellular-resolution imaging in organotypic cultures of the adult zebra finch brain, including portions of the intact song circuit. These *in vitro* networks can survive for weeks, and display mature neuron morphologies. Neurons within the organotypic slices exhibit a diversity of spontaneous and pharmacologically induced activity that can be easily monitored using the genetically encoded calcium indicator GCaMP6. In this study, we primarily focus on the classic song sequence generator HVC and the surrounding areas. We describe proof of concept experiments including physiological, optical, and pharmacological manipulation of these exposed networks. This method may allow the cellular rules underlying sparse, stereotyped neural sequencing to be examined with new degrees of experimental control.

**Highlights:**

- *Organotypic brain slices from adult zebra finch (Taeniopygia guttata), expressing the calcium indicator GCaMP6, can be cultured and maintained for at least several weeks and display spontaneous and evoked calcium transients.*

## INTRODUCTION

Sequential patterns of neural activity, distributed across functionally distinct brain regions, underlie a variety of behaviors including movement (Pastalkova et al. 2008), memory (Robinson et al. 2017; Cai et al. 2016; Robinson et al. 2017), navigation (Harvey et al. 2012; Morcos & Harvey 2016; Harvey et al. 2012), and sensation (Vinje & Gallant 2000; Olshausen & Field 2004). Dysfunction in the spatio-temporal patterns of activity in these networks are implicated in a variety of human disease including Parkinson’s (Stahl et al. 2009; Daviaud et al. 2014; Duff et al. 2002) and Alzheimer’s (Christian Humpel 2015; Mewes et al. 2012). It is still not known how connectivity of these networks is established, or how spatio-temporal patterns of activity are maintained, modified, and coordinated within these networks (Peters et al. 2017; Gulati et al. 2017). Rapid progress is being made in studies of sequence production in songbirds, but methods to investigate the homeostatic mechanisms underlying sparse firing networks are lacking (Picardo et al. 2016; Vallentin et al. 2016; Kosche et al. 2015; Markowitz et al. 2015). We reasoned that if organotypic cultures made from thick sagittal sections, that contained large portions of the song system, could produce sparse and possibly even stereotyped activity *in vitro,* this would provide a basis to examine the cellular processing underlying homeostatic control of cells.

Neuron cultures have already yielded great insights into homeostatic plasticity mechanisms, such as Long-Term Potentiation, LTP, Long-Term Depression, LTD (Karmarkar et al. 2002; (Turrigiano et al. 1998; Turrigiano 2008), synaptic scaling, structural synaptic plasticity (Yasumatsu et al. 2008; Tominaga et al. 1994), self-organization of neural networks (Johnson & Buonomano 2007) (Cruz-Martín & Schweizer 2008), and the underlying mechanisms of brain disease (Glykys & Staley 2015). Long term organotypic cultures provide a unique system where emergent properties of neural networks can be studied at the cellular and molecular level, with extremely precise experimental control. These exposed networks are ideally suited for gene transfer techniques using electroporation and synthetic/modified viruses to express genetically encoded proteins for optical recording and control (Andersson et al. 2016; (Rathenberg et al. 2003), repeated multi-electrophysiological recordings and stimulation (Dong & Buonomano 2005), and subsequent whole-circuit reconstruction using electron microscopy (Buchs & Muller 1996).

A diverse array of protocols for organotypic culture systems exist (C. Humpel 2015). The common preparation involves a stable culture medium and sufficient oxygenation and incubation. Under these conditions, nerve cells often develop a tissue organization that closely resembles that observed *in situ* (Gähwiler 1997). Organotypic cultures can allow aspects of structural and synaptic organization of the original tissue to be preserved, and are ideally suited for studying and precisely manipulating the interplay between locally and distally generated activity patterns, which is not yet possible *in vivo*. The feasibility of studying neuronal pathway development in long-term explant cultures has been shown across many mammalian neural systems (Dailey & Smith 1996; Molnár & Blakemore 1999; Toran-Allerand 1991).

However, there are clear limitations to mammalian organotypic culture as a model of circuit function. Although cultures typically maintain their local connections in slice, they are cut off from many, if not all of their downstream and upstream projection targets. Axotomy and oxygen deprivation in the interior of the culture causes widespread necrosis and apoptosis in the early phases of organotypic brain slice (C. Humpel 2015). The rapid population decrease can lead to the collapse of the well-defined local laminar cytoarchitecture in cortical structures; an organization that is thought to be key for supporting computations. Hippocampal networks have evolved as the predominant *in vitro* cultured systems in part because they can partially retain the cytoarchitecture of the tissue of origin (Gähwiler 1981). While these cultures can be coerced to recapitulate *in vivo* like activity, organotypic slice cultures are seizure prone (Bausch 2009). This may be the result of less naturalistic external drive from sensory regions that do not typically survive the culture process or of an increase in the probability of synaptic connectivity (Pavlidis and Madison, 1999; Cruz-Martín & Schweizer 2008). In addition, it is not trivial to maintain long-term cultures of adult mammalian tissue, although there have been successes in some adult slices with tissue-specific culture conditions optimized for adult organotypic slices (Lossi et al. 2010; Wilhelmi et al. 2002; Kim et al. 2013).

The nucleated cytoarchitecture of the avian brain may provide advantages over mammalian organotypic brain slices. Under typical culture conditions, distant, functionally related brain areas can be cultured simultaneously, and the nuclear organization of the slices is preserved for weeks in culture, with protein synthesis continuing at an essentially constant rate for weeks (Holloway & Clayton 2001). A nuclear organization may also be ideal for preserving local anatomical connections since heterogeneous populations of cells self-organize within nuclei with relatively little or no imposed topographical organization within the nucleus (Jun & Jin 2007; Johnson et al. 1995). This local connectivity is presumably driven by local genetic and molecular cues, which are easily recapitulated in organotypic culture (Holloway & Clayton 2001). In addition, the songbird brain has well-described regenerative properties (Kirn et al. 1994; Alvarez-Buylla et al. 1992), where newborn neurons migrate and can repair the adult brain (Kubikova et al. 2014). Many song-motor related nuclei in seasonal breeding songbirds are known to oscillate greatly in size and cell number from one season to the next, while the stereotypy of song can remain remarkably stable for years (Kirn et al. 1994; Alvarez-Buylla et al. 1992)

For our purposes, the songbirds offer an even greater benefit. Zebra finches in particular are excellent model for studying neural sequence generation. Among songbirds, the song circuit of zebra finch is one of the most studied and one of the best understood. These animals produce an extremely robust and stereotyped motor behavior that is known to be driven by an ultra-sparse code in HVC, the songbird analogue of mammalian premotor cortex and presumed neural sequence generator (Hahnloser et al. 2002). While temporally-ordered neural activity has also been observed in other species in the context of various sequential behaviors (Harvey et al. 2012; Pastalkova et al. 2008; MacDonald et al. 2011), the extreme precision and sparsity of the songbird premotor projection cells in HVC are unmatched. Sparsely active projection neurons in HVC are precisely time locked to song, and are thought to generate a timing signal (Hahnloser et al. 2002). One of the most prominent theoretical models of neural sequence generation is the synfire chain(Long et al. 2010; Abeles 1982; Li & Greenside 2006), in which pulses of synchronized spiking activity propagate robustly along a chain of cells connected by highly redundant, effectively feedforward excitation. In the songbird, there is evidence of a spatially recurrent excitatory chain, passing repeatedly through zones of local feedback inhibition (Markowitz et al. 2015; Cannon et al. 2015). How is this spike timing precision maintained in the presence of biological noise? What makes neural sequence generation in this context robust, and how does it evolve over time (Clopath et al. 2017; Peters et al. 2017; Chambers & Rumpel 2017; Lütcke et al. 2013)?

Here, we present proof of concept calcium imaging data from long-term organotypic cultures of the song system using genetically encoded indicators. In this preparation, we describe tools for monitoring and manipulating the physiological and pharmacological properties of these neuronal circuits *in vitro,* providing new tools to examine the cellular and homeostatic mechanisms of sparse sequence generation.

## METHODS

### Subjects

All procedures were approved by the Institutional Animal Care and Use Committee (IACUC) of Boston University. Zebra finches (*Taeniopygia guttata*), were raised by their parents in our lab aviary or purchased as adults from a supplier. All animals were kept on a 14 h light-dark cycle. Organotypic slice data used in this work were collected from four adult male zebra finches (>120 days post-hatch). One finch provided the *in vivo* data used in **Figure 1a.**

**Fig 1.**
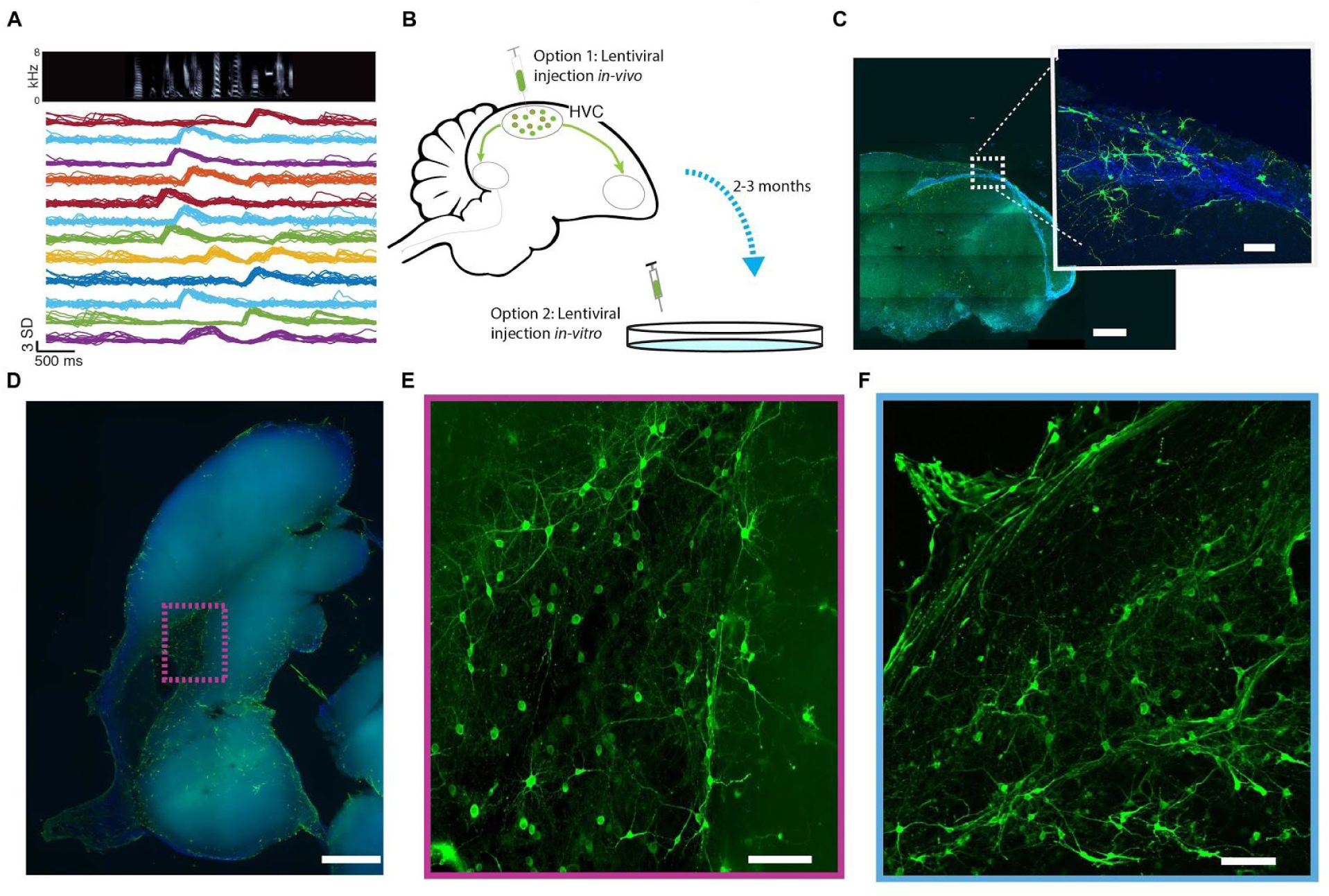
Neurons survive in organotypic cultures of the songbird brain. **(A)** Wide-field recordings of calcium activity in HVC of the awake behaving songbird show sparse, sequential activity, that is stereotyped across a single day. *Top:* Audio spectrogram of a bird’s song. *Bottom:* Peak ΔF/F^0^ -normalized, trial-averaged activity from 16 song motif-locked neurons during the song in *top*, from 20 song-aligned trials during a single day of imaging. *Y-axis* is in standard deviation (SD). **(B)** Adult songbird culture overview, where lentivirus containing RSV-GCaMP6 was injected into HVC. Alternatively, virus was bulk applied onto the culted slice at 3 and or 8 DIV. **(C)** Organotypic brain slice cultured from adult birds shown after 2-3 weeks in culture. Scale bar indicates 1mm. *Inset:* Area of neurons centered in the ventricle below HVC. Scale bar indicates 100μm **(D)** RSV-GCaMP6s was applied on 3&8 DIV, and the slices were fixed on 34 DIV (N=3). Immunostaining for anti-GFP (*green*) showed GCaMP6s infected cells after 5 weeks in culture DAPI counterstain (blue) for nuclei. Scale bar indicates 1mm **(E)** inset from **D.** Scale bar indicates 1mm. **(F)** A different slice, showing cells expressing GCAMP6f (30 DIV). Scale bar indicates 100μm.

### Organotypic brain slice (OBS) cultures

Animals were perfused. A total of 40 slices from four animals were cultured. Of these, 25 slices were imaged in single session time-lapse recordings ranging in duration from 1-4 hours. 15 of these slices yielded data suitable for analysis. Of these imaged slices, four were infected *in vivo*, the remaining slices (n = 11) were infected *in vitro*. A subset of these slices were fixed and used for histology plating on 15 or 30 days *in vitro* (DIV). Three additional slices (34 DIV) were fixed without imaging for histology. Slices were not cultured beyond 48 DIV. See **Supplemental Table 1** for a full description of slices and conditions used in this experiment.

Slice cultures were prepared according to a modified protocol as previously described (Stoppini et al., 1991; De Simoni and Yu, 2006). Briefly, before animal decapitation, the membrane of insert (Millipore: PICM03050) was placed in separate well of a six-well culture plate (Nunc six-well dishes: 140675) with 1 ml culture medium underneath (per 100ml: 50 ml MEM with Glutamax-1 (Gibco-Invitrogen: 42360-024), 23 ml EBSS (Thermo Fisher Scientific: 24010043), 6.5 mg/ml D-glucose (Sigma: G8270), 40-50u/ml penicillin–streptomycin (Life Tech: 15140-122), 25 ml horse serum and 0.06 ml Nystatin (Sigma: N1638)), and a sterile confetti (Millipore: FHLC01300) was placed on the membrane of insert. The six-well culture plate was then held at 37°C in a 5% CO^2^ incubator for at least two hours but not more than 24 hours before plating. The birds were anesthetized with pentobarbital (250mg/kg, IM). Following decapitation, the brain was removed and placed in ice-cold slicing buffer (SB) which contains modified Earle’s Balanced salt solution (Ca^2+^ free, Mg^2+^-free, phenol red free; Thermo Fisher Scientific: 14155063) plus 25mM HEPES. From slicing until plating, all equipment and solutions were kept ice cold. Brains were first bisected at the midline, then glued (Loctite) to the vibratome stage with the cut surface up, and flooded with SB. Parasagittal slices (180-300um) were cut using a Leica vibratome (Leica Biosystems Inc: VT 1000s). This yielded 4(± 2) intact slices per bird containing visible portions of both HVC and HP. The slices were then transferred to sterile centrifuge tubes of ice-cold SB using a sterile 3 ml modified plastic pipette (wider opening) and stored until plating. The slices were then incubated at 37°C in a 5% CO^2^ incubator. 1.0 ml fresh culture medium was replaced after the first hour *in vitro* and then again every 24 hours. This exchange was repeated every other day for the duration of the culture. Slices could be maintained and used for 4-5 weeks after plating. We determined empirically that viable OBS slices could be maintained for up to 5 weeks after plating. OBS were regularly inspected for signs of necrosis or infection. Beginning at one week after plating days *in vitro* (DIV). By 7 DIV, slices had adhered and could be used for successful imaging.

### Imaging Media Preparation

Membranous OBS were placed in the (pre-warmed) chamber at least 30 minutes before imaging and kept at 34°C throughout the imaging period (up to 4 hours). For imaging, the membrane with the cultured slice was removed from the incubator and transferred to the imaging microscope in heated, fresh 95% O^2^ /5% CO^2^ aerated artificial cerebrospinal fluid (ACSF) containing:

> **Solution 1:** 124 mM NaCl, 26 mM NaHCO^3^, 3 mM KCl, 1.25 mM NaH^2^ PO^4^, 1.5 mM MgCl^2^, 2 mM CaCl^2^, and 20.0 mM glucose (Gao et al. 2017), or:
>
> **Solution 2:** 126 mM NaCl, 6 mM NaHCO^3^, 3 mM KCl, 1.25mM NaH^2^ PO^4^,2 mM MgSO^4^, 2 mM CaCl^2^, and 10.0 mM glucose (Long et al. 2010).

The membrane with the slice was then placed in an open perfusion chamber (Harvard Apparatus:RC-26G) slice-side up and was held in place by a titanium harp with three single nylon fibers glued on. The harp provided enough pressure to hold the slice in place. The perfusion chamber was gravity fed with aerated, heated recording solution. Slices were fully immersed in continually flowing recording solution with a perfusion rate was around 2.5-4 ml/min, similar to what has been described by others (Dailey and Smith, 1996). Temperature was maintained and monitored with a dual automatic temperature controller (TC344C, Warner Instruments). In some experiments a KCL solution was used to evoked activity in OBS, because the presence of extracellular potassium can stimulate calcium influx in neurons (Willeumier et al. 2006). For KCl depolarization, OBS were slowly perfused with recording solution containing 10, 20, 40 mM KCl for 5-10min, followed by washing with normal recording solution. We estimated this media exchange took about one minute.

### Virus production

To monitor the activity of organotypic slice neurons we used plasmids pGP-CMV-GCaMP6s and pGP-CMV-GCaMP6f (Addgene # 40753 and #40755, respectively). These plasmids were a gift from Douglas Kim laboratory. pHAGE-RSV-GCaMP6s, pHAGE-RSV-GCaMP6f, and pHAGE-RSV-GFP plasmid maps and sequences can be downloaded from the vectors page of the laboratory of Darrell Kotton (www.kottonlab.com). These viruses were previously described and used by our lab (Liberti et al. 2016). The viruses were packaged in HEK 293T cells and titered on FG293 cells, with titers ranging between 1.2-2.3 x 10^10 infectious particles/mL. Specific plasmids can be found and ordered on addgene: https://www.addgene.org/Darrell_Kotton/

### Viral knock in of reporter of neural activity

To label neuronal cells in OBS cultures we performed viral injections either *in vitro* or *in vivo*.

#### In-vivo preparation

For injections of lentivirus expressing GCaMP6s under the Rous sarcoma virus (RSV) (pHAGE-RSV-GCaMP6s, titer, 1.2-2.3 x 10^10 infectious particles/mL), as previously described (Liberti et al. 2016; Markowitz et al. 2015) in brief, a small burr hole was made in the cranial surface over HVC, and a small glass pipette was lowered 150um from the pial surface. 250 nl of virus was inoculated into the brain in up to four injection sites. To guide the injection of virus, the bounds of HVC were determined through fluorescence imaging of a DiI retrograde tracer which was injected in Area X one week prior.

#### In-vitro viral application

OBS containing HVC was infected with pHAGE-RSV-GCaMP6s. Infection was performed at 4-9 DIV by gently lowering a micropipette containing 10μL of 1:100 diluted viral stock solution (original titer, 1.2-2.3 x 10^10 infectious particles/mL) into the target location of the tissue and applying positive pressure. Experiments were performed 6-25 days after viral transduction. Due to the extremely low baseline resting fluorescence of GCaMP6, in some instances OBS were co-cultured with a diluted RSV-GFP in order to find and focus on areas with potential GCaMP6s expression. We excluded cells co-infected with GFP and GCaMP6s from calcium imaging data analysis because the spectral overlap of the fluorescent signals.

### Calcium Imaging

#### Calcium Imaging in Vivo

Calcium imaging the activity of HVC excitatory projection neurons during singing was performed as previously described (Liberti et al. 2016; Liberti et al. 2017)

#### Calcium imaging acquisition in vitro

GCaMP6 imaging was performed with two different detection systems attached to the same custom microscope. Wide-field calcium imaging was performed using an Olympus BX observation tube, eyepieces and fluorescent filter cube housing, custom LED for excitation, and sCMOS camera (OrcaFLASH4.0, Hamamatsu, Bridgewater, NJ). Two-photon calcium imaging was performed using a stock movable objective microscope (M.O.M Sutter Instruments). Excitation was provided by a Tsunami titanium: sapphire laser pumped by a 5W Millennia pump laser (Spectra-Physics, Santa Clara, CA) The output was mode-locked with the aid of a Model 3955 Mode-locker(Spectra-Physics). The ultra-short pulses were 150fs wide and had a repetition rate of 80MHz. The beam was spectrally profiled with a compact spectrometer (Part #CCS175 ThorLabs, Newton, NJ) and software (Splicco 4.3.73, Thor) and tuned to 910 nm. In order to minimally damage the slices, power was attenuated to the lowest usable level by means of a Model 302RM amplifier and model 350-80LA Electro-Optic Modulator (Conoptics, Danbury, CT). Upon exiting the Pockels cell, the beam passed through steering optics, then past a shutter, (ThorLabs SC10 shutter) directly onto the periscope of the resonant scanner (RESSCAN, Sutter Instruments). Excitation was delivered to (and collected from) the specimen via a Nikon LWD 16x /0.8W water immersion lens. Fast Z scanning was accomplished with a PFM 450 piezo objective drive (ThorLabs). Detection of fluorescence was by a Hamamatsu wide field of view photon counting photomultiplier (H10769-40). Sequential recordings were made at 15.2 frames per second of a 512x512 field of view with a zoom factor of 3. A Sutter manipulator (ROE 200, Sutter Instruments, Novato, CA) was was used for controlling the x, y and slow z of the stage. A Lenovo tower computer with an Intel i7 4790 CPU operating at 3.6Ghz and 32Gb of RAM was used for all acquisition and analysis. MATLAB R2015B (MathWorks, Natick MA) and ScanImage 2015 (Vidrio Technologies, Ashburn, VA) were used to interface with the hardware. The resonant scanner was operated at a frequency of 7910 Hz.

#### Analysis of Calcium imaging data

In the GFP recording channel, individual cell somata were selected as the Region of interest (ROI) and the mean intensities in the ROIs in each frame were determined. Raw data delivered in the form of a linear 16-bit intensity scale were first plotted as fluorescence intensity versus time. The background fluorescence measured near a ROI was then subtracted from these raw data. The F^0^ (resting fluorescence) image was generated by averaging the fluorescence intensities of 10–50 frames from the image stack in a time window when there was no significant change in calcium fluorescence. Subsequently, data were normalized to the mean fluorescence intensities [ΔF/F_0_ = (F − F_0_)/F_0_], allowing the comparison of data across experiments. At the end of the imaging session, some slices were immediately fixed in 4% paraformaldehyde for further analysis.

#### Organotypic slice fluorescent immunohistochemistry

At the end of the experiment, the slices were fixed overnight by 4% paraformaldehyde in 0.1 M PBS and were used for immunohistochemical staining of GFP and DAPI. Non protein binding was blocked with 10% normal donkey serum. The primary antibody used to detect GCaMP6s was a rabbit Anti-GFP (1:500, NB600308, Fisher Scientific). Following primary incubation for overnight at 4°C and of PBS washing (3X), Alexa Fluor® 488 AffiniPure Donkey Anti-Rabbit IgG (H+L) (1:500, 711-545-152, Jackson ImmunoResearch Laboratories) was applied. The slices were coated with mounting medium containing 4’,6 diamidino-2 phenylindole (DAPI, VECTASHIELD). OBS were visualized and the images were captured using a FV10i confocal microscope (Olympus) with Olympus FV10i software and Nikon Ni-E Motorized Microscope for Fluorescence with Monochrome and Color Cameras.

## RESULTS

### Projection neurons survive in-vitro explant culture, and retain neural specific morphology

Projection neurons *in vivo* produce ultra sparse, motif-locked patterns of neural activity, and these can be recorded *in vivo* using head-mounted microscopes (**Figure 1a).** (Markowitz et al. 2015; Liberti et al. 2016) While these cells produce robust, reliable sequences in the adult *in vivo*, we asked whether these neurons would spontaneously recapitulate similar patterns in a cultured explant. To determine if cells would remain viable after the explant procedure, we removed the brain from adult Zebra finch, where HVC had been infected with the calcium indicator GCaMP6 and the cells had been identified as producing time locked sequences *in vivo*, **Figure 1a.**

A subset of slices received additional virus applications to infect multiple portions of the song **system** beyond HVC **Figure 1b** (see **Supplemental Table 1**) and some slices only received a bulk application of virus directly onto the slice. This method labeled far more cells with GCaMP6s, but the identity of these cells could not be determined **Figure 1b**. Both *in vitro* and *in vivo* viral delivery yielded obvious GCaMP6 infected cells, some of which were responsive. Organotypic slices were cultured from adult birds, and cells survived even after several weeks **Figure 1c-f**. To examine cell morphology, several slices were fixed (34 DIV) and immunostained with an anti-GFP antibody to label the GCaMP6s infected cells after 5 weeks in culture (n=3) **Figure 1d-e**. Numerous sections contained GFP-positive cell networks exhibiting a variety of morphologies **Figure 1e-f**.

### Pharmaceutical access to zebra finch song system organotypic culture

Imaging slices in a standard perfusion chamber perfused with oxygenated recording solutions, we asked if basic pharmacological techniques known to work acutely *in vivo* could be used to manipulate neurons in slice. After monitoring a subset of slices showing no obvious activity after periods of 60 minutes, slices were stimulated with KCl. Elevated external KCl can depolarize neurons and stimulate calcium influx (Fairless et al. 2013). These resulted in robust activation of neurons **Figure 2.** Once an area was identified, calcium activity was recorded in response to high dose (above 30 mM) of KCl stimulation (**Figure 2a-c,** also see **Supplementary Video 1**). For KCl depolarization, slice s were treated with Recording ACSF (RCSF) containing 15 mM KCl for 1-2mins, followed by washing with normal RCSF with KCl 3mM 10 minutes before treating with KCL again. The low dose KCl and washout can reversibly activate GCaMP6f cells **Figure. 2d**. Many networks showed extremely sparse activation, and in our hands KCl application was a beneficial way to identify regions of infected cells. KCl depolarization is reversible: slices were treated with RCSF containing 15 mM KCl for 1-2mins, followed by normal RCSF with KCl 3 mM washing 5 mins and 30 mM KCL was treated again. The low dose KCl and washout can reversibly activate GCaMP6f cells **Figure 2d**.

**Fig 2.**
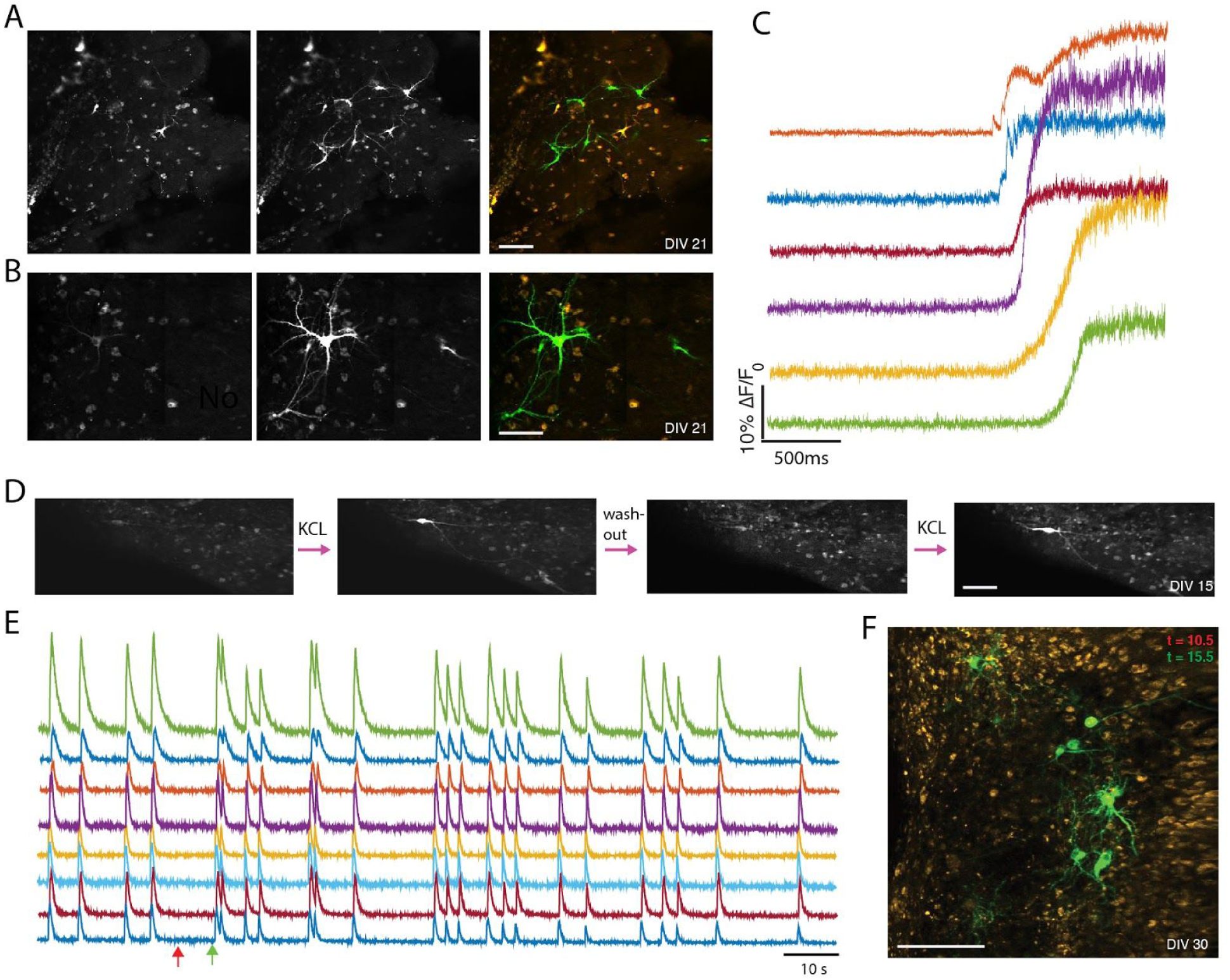
Pharmaceutical manipulation of Organotypic brain slice infected with RSV-GCaMP6f. **(A)** Calcium activity in response to KCl stimulation. *Left*: baseline fluorescence. *Middle*: evoked calcium signal after the application of KCl. *Right:* overlay, red = before, green = after. **(B)** same as **A**, with a different slice. (**C)** Examples of calcium activity from ROIs shown in **B** 1-2 minutes after a high dose of KCl. **(D)** A low dose KCl and washout can reversibly activate GCaMP6f cells. Two sequential periods of KCL application are 1 minute and the washout time is 5mins. Scale bars =100μm. **(E)** ΔF/F^0^ traces from multiple ROIs, as an example of synchronous network activity in the presence of GABA-ergic antagonism (30 DIV, 37°C, see methods). Red and green arrows mark time points used in the next panel. **(F)** Overlay of an ‘active’ period (green) and a ‘quiet’ period (in red), with corresponding colored time marks indicated in **E.** Five seconds elapsed between green image and red images. All scale bars =100μm.

### Recording spontaneous network activity in songbird organotypic brain slice

Finally, we asked whether network activity can be recorded spontaneously from these explants. Neurons develop synaptically driven spontaneous bioelectric activity in mammalian thalamocortical organotypic slice (Klostermann & Wahle 1999). Songbird organotypic slice exhibited asynchronous single cell calcium spikes by at least 15 DIV (see **Supplementary Video 2**) and as well spontaneous, somewhat asynchronous calcium waves by 30 DIV **Figure 3 a-b** (also see **Supplementary Video 3**). The Organotypic brain slice could be induced to exhibit a more uniform synchronous calcium waves at 30 DIV using an antagonist, Picrotoxin (PTX) to block the GABAA receptors and drive an overall increase in activity (**Figure. 2e-f)**.

**Fig 3.**
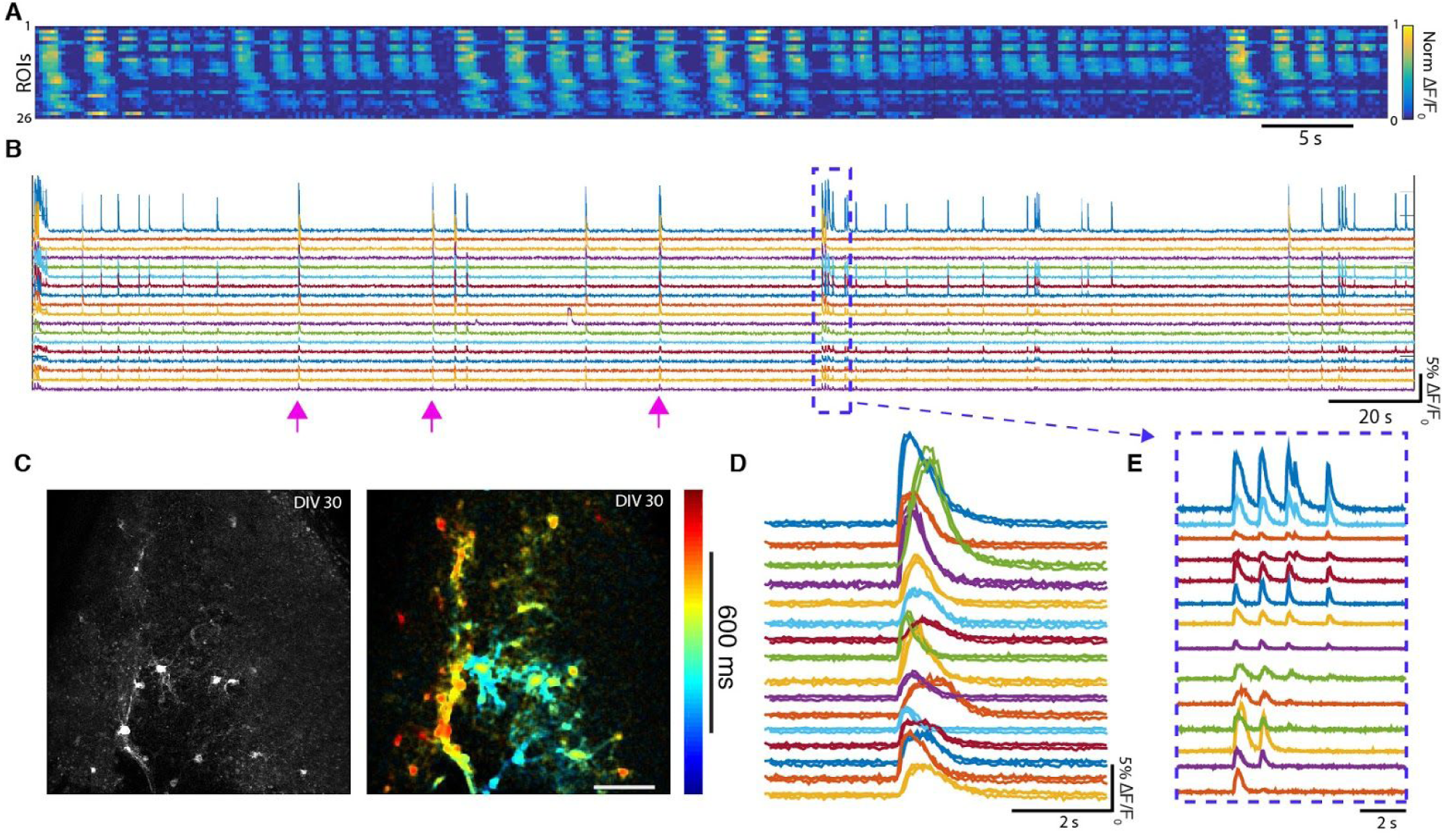
Spontaneous bursting activity exhibited in songbird organotypic brain slice recorded at 33-34 degrees Celsius. (**A-B)** Calcium transients observed in spontaneously active cells. Each row is one cell and the gaps between calcium bursts have been edited out, revealing the stereotypy of the burst event. **(B)** Calcium transients shown in **A** without the silences edited out. Each color is a different ROI. **(C)** *Left:* maximum projection of cells shown in **A-B.** *Right:* image from **C** false colored by the timing of max pixel intensity reveals a wave travelling from blue to red. **(D)** Pink arrows in **B** mark three separate bursts of spikes that were overlaid in this panel. Bursts contain highly stereotyped activity across cells. **(E)** Another event showing a gradual tapering in cell participation in a rhythmic burst.

### Spontaneous and asynchronous network activity in cultured HVC networks

In addition to highly correlated synchronous activity, we also observe sparse, slow sequential activation of cells in normal culture media. However, it is important to note that these sequences were over an order of magnitude slower that what is typically seen in vivo (Markowitz et al. 2015; Liberti et al. 2016). We hypothesized that our culture media may not be ideal for revealing naturalistic functional activity, so we made slight modifications of the culture media, using 50% Ca^2+^/Mg^2+^modified recording solution 2 (described in detail in the methods). This revealed a rich diversity of asynchronous network activity, which was observed for 2 hours with different slow and fast calcium transients in different cells (**Figure 4a)** or the same cell (**Figure 4d)**.

**Fig 4.**
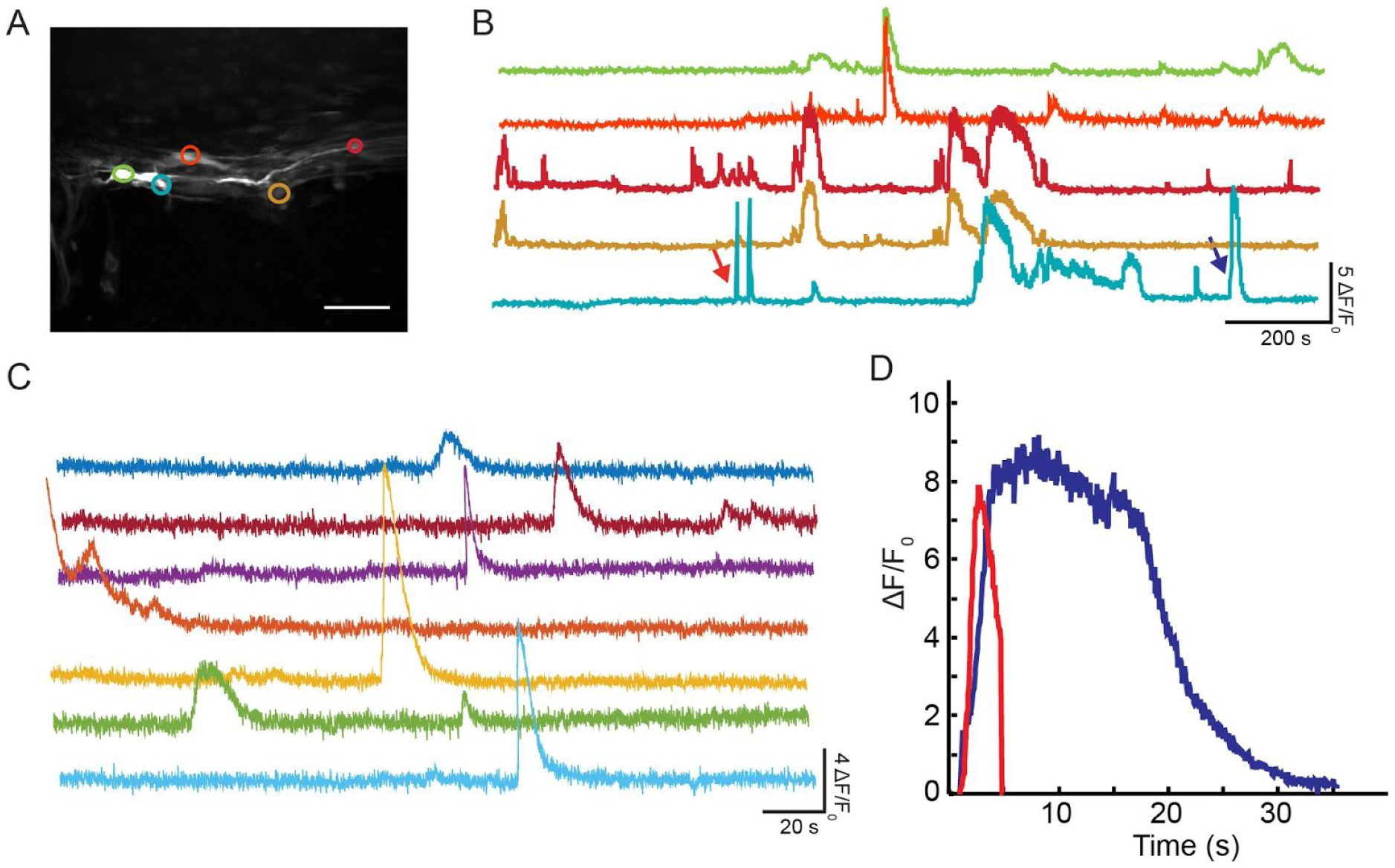
Organotypic brain slices also exhibit slow spontaneous, asynchronous network activity. **(A)** Maximum projection of a slice exhibiting spontaneous activity and their corresponding traces, color coded. Scale bar indicates 50μm. **(B)**. Recordings were performed with 30 DIV (34°C in half concentration Ca^2+^/Mg^2+^modified solution 1). (**C)** In another slice (30 DIV, 34°C, solution 1), sparse asynchronous activity could be observed. However, these sparse activations are significantly slower than transients observed in-vivo (see **Figure 1a**) and did not repeat over the time we recorded. **(D)** ROIs showed a variety of slow timescale activation. In one ROI example, showing a short transient (*red trace*) early in the trial and a broader response (*blue trace*) later in the session.

## DISCUSSION

In this preliminary work, we demonstrate that cells can survive in songbird organotypic slice culture for many weeks, and that these cells appear to be functional. We provide several examples of cells in slices containing the partially intact song system that appear to reproduce some of the diversity of the anatomical and functional properties of their corresponding circuits *in vivo*. These results provide a compelling argument to further develop organotypic culture as a method to study the functional, genetic, neurochemical, and anatomical basis of sparse neural sequence generators *in vitro*. Multi-month songbird organotypic lines could yield critical insight into the underlying principles involved in the assembly and function of neuronal circuits, including homeostatic plasticity (Turrigiano et al. 1998; Turrigiano 2008) and repair in response to neurodegeneration. Avian brains are particularly well suited to the study of network reorganization and repair, in light of well documented ability for the songbird brain to undergo extensive circuit repair after damage (Kubikova et al. 2014) and coupled with seasonal cycles of neurodegeneration and neurogenesis in the adult that constantly destroy and rebuild circuits that produce the same behavior (Kirn et al. 1994; Alvarez-Buylla et al. 1992). In addition, it is possible that nuclei can operate sufficiently even when intra-regional connections are completely severed, suggesting that in principle, even thick sections that only contain partial nuclei could maintain a complete, functional, distributed network (see **Supplemental Figure 1)** (Poole et al. 2012; Hamaguchi et al. 2016; Roberts et al. 2017).

Advances in techniques to monitor and perturb neural systems are maturing rapidly and have been demonstrated viable in the songbird, and should directly translate to organotypic culture of the same tissue (Roberts et al. 2012; Liberti et al. 2016). Replacing blood with oxygenated media decreases light scattering, making these cultures ideal for high-resolution structural and functional imaging across the entire exposed tissue volume (Ji et al. 2016). Alternatively, it is feasible to culture brain tissue on a mesh of electrodes, providing high-throughput electrical stimulation and recording (Liu et al. 2015; Zhou et al. 2017). Viral approaches to knock down or overexpress genes, silence or activate neurons, or render neurons responsive to light or selective drugs will permit cell-type specific, high-temporal and spatially precise all-optical physiology across entire functional networks.

This study contains a significant number of unanswered questions. The extent to which cells maintained and generated within songbird organotypic culture recapitulate the cellular and genetic diversity and circuit functionality of their *in vivo* counterparts is not addressed in this study. Future experiments may rely on genetic targeting strategies to identify and manipulate defined cell types. In addition, while a wide diversity of activity was seen in our organotypic cultures, the timescale of this activity is almost 10x slower than what is seen *in vivo* in the corresponding brain regions. The baseline spontaneous activity was low, as it is in many song circuits *in vivo* in absence of motor production (Mooney 2000; Kozhevnikov & Fee 2007), but it is unclear if this is abnormal for these circuits. This preliminary study was not able to fully characterize the optimal conditions for evoking spontaneous activity. In all cases, it is unclear if the cells that produced activity were specifically derived from the *in vivo* song circuit or from cells from nearby regions. Many protocol deviations were attempted and as a result, observations of network activity were often limited to a few examples. A substantial effort will be needed to validate if networks that arise in songbird neural culture match their *in vivo* counterparts. Future directions should include characterization of projections patterns of specific networks using viral retrograde methods (Cruz-Martín et al. 2014) in combination with chronic monitoring of activity using fluorescent indicators. These methods should allow for the characterization of specific networks within the avian organotypic brain slice preparation.

However, for many experiments, the exact characterization of songbird organotypic culture is unimportant. The integration of real-time optical and electrical control and monitoring opens up fascinating opportunities. Songbird neural culture could provide an ideal physical substrate for a ‘biological computer’ where operations are programed by electrically or optically stimulating select cells to write-enable, or entrain sequences, to enforce certain connections between cells, or to drive particular patterns of protein or nucleotide expression. Real-time feedback on patterns of activity could not only discover circuit plasticity rules, but it could also provide a testbed for implementing biologically inspired machine-learning or reinforcement learning paradigms (Clancy et al. 2014; Ganguly et al. 2011; Morcos & Harvey 2016). Modulation of neural activity could shape and store information in network patterns, and these populations could be ‘programmed’ to process stored data. Culturing can be automated, and many cultured networks can be linked together as specialized functional components of a multi-layer system: a biological computational engine. Potentially, we can begin to observe the details of how natural neural network self-organization leads to emergent solutions to computational problems.

## SUPPLEMENTARY MATERIAL

**Supplemental Figure 1:**
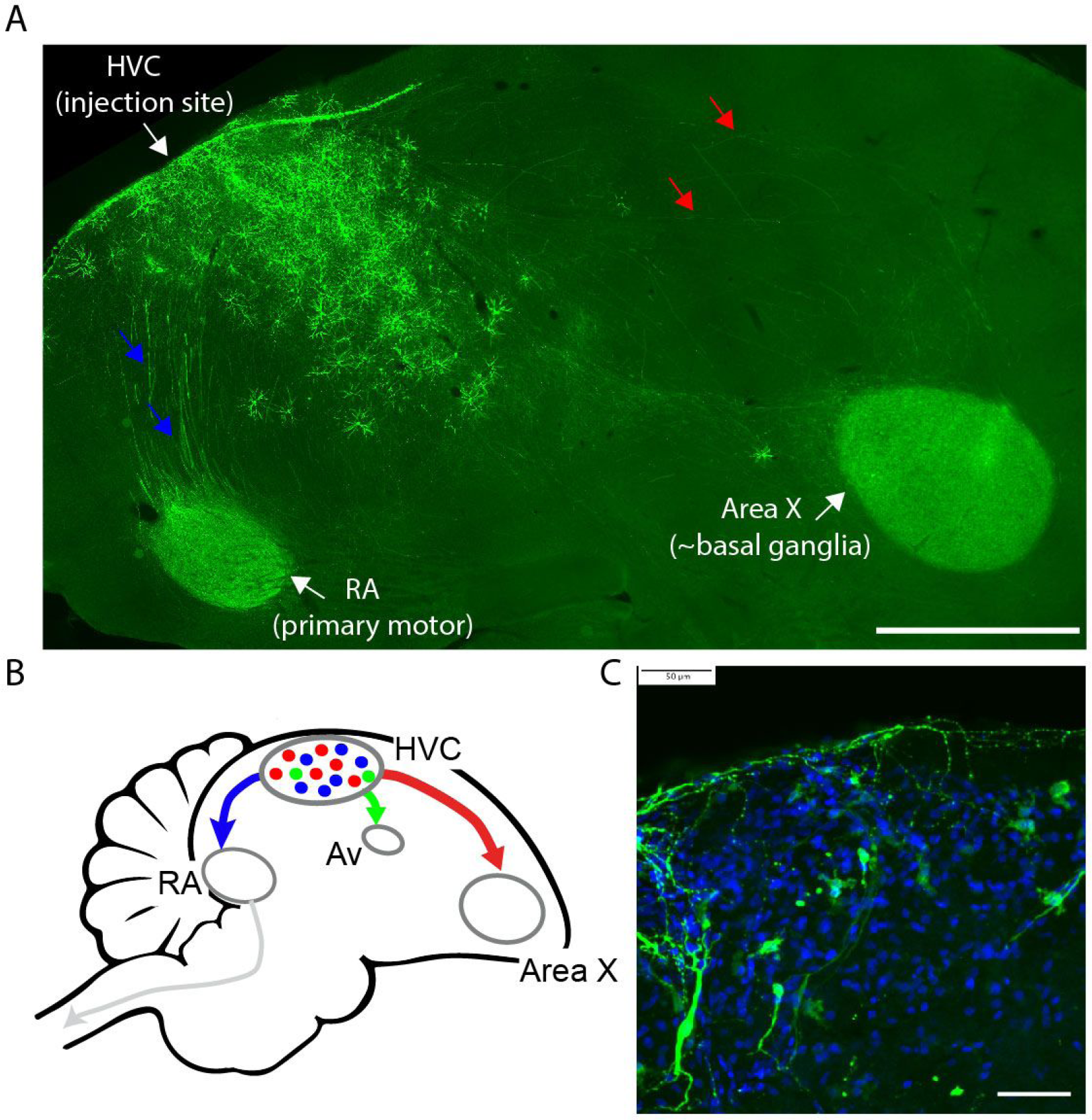
Sagittal brain slices can contain major portions of the song circuit, including projection targets. **(A)** Acute histological section of the zebra finch brain, with an injection in HVC. *Red* and *blue* arrows indicate axons meandering through the thin section. Axons aggregate in downstream nuclei Area X and RA- and the axonal path of many of these projections are contained in this slice. Green = GFP stain. Scale bar indicates 500μm **(B)** Schematic of the songbird brain, showing projection targets from HVC. **(C)** Many cells survive the slice process. *Blue* = DAPI, *Green* = GFP. Scale bar indicates 50μm.

**Supplemental Table 1:** Summary of the conditions for organotypic slices from all experiments used in this study. Columns from left to right: Animal of origin, Days *in vitro* (DIV) at the time of imaging or histology, Temperature of media during imaging, method viral application, Imaging culture media (see **Methods**), and Summary of pharmaceutical manipulation. This table does not include the animal used in **Figure 1a**, or the acute histology from **Supplemental Figure 1.**

| Animal | DIV | Temp (°C) | Virus application | Media | Summary |
| --- | --- | --- | --- | --- | --- |
| LOR25RB | 28 | 28 | *in vivo* | 1 | Spontaneous Activity |
| LNO26 | 10 | 27.2 | *in vivo* | 1 | Spontaneous Activity |
| LNO26 | 21 | 28 | *in vivo* | 1 | KCL 50-120mM |
| LNO25 | 13 | 28 | 9 days *in vitro* | 1 | KCL 40mM |
| LNO25 | 21 | 27.4 | 9 days *in vitro* | 1 | KCL 40mM |
| LNO25 | 21 | 27.5 | 9 days *in vitro* | 1 | KCL 20-40mM |
| LNO25 | 29 | 28.8 | 9 days *in vitro* | 1 | Ca:Mg=2:1; PTX 100μM;<br>KCL 20mM |
| LNO25 | 2 | 33-41 | *in vivo* | 2 | PTX 10μM |
| LNY40 | 8 | 33 | 3 days *in vitro* | 2 | Spontaneous Activity |
| LNY40 | 30 | 32.5-36 | 3 & 8 days *in vitro* | 1 | PTX 2.5-10μM KCL<br>20mM |
| LNY40 | 30 | 33 | 3 days *in vitro* | 1 | Spontaneous Activity |
| LNY40 | 15 | 33-35.8 | 3 & 8 days *in vitro* | 1 | Spontaneous Activity |
| LNY40 | 15 | 32.5 | 3 & 8 days *in vitro* | 1 | KCL 5nM-30mM |
| LNY40 | 15 | 32.6 | 3 & 8 days *in vitro* | 2 | Ca:Mg=4:3 (50% less than<br>normal) |
| LNY40 | 30 | 33-34 | 3 & 8 days *in vitro* | 2 | Spontaneous Activity |
| LNY40 | 15 | N/A | 9 days *in vitro* | N/A | Histology only |
| LNY40 | 34 | N/A | 3 & 8 days *in vitro* | N/A | Histology only |
| LNY40 | 30 | N/A | 3 & 8 days *in vitro* | N/A | Histology only |

***Supplementary Video 1:*** Cells active in response to high dose (above 30 mM) of KCL stimulation (21 DIV). Movie playback speed is increased by a factor of 10, and temporally smoothed by a factor of 4. Link: https://youtu.be/_0DuzXjOCGU

***Supplementary Video 2:*** Songbird organotypic slice exhibiting asynchronous single cell calcium activity (30 DIV). Movie speed is increased by a factor of 10, and temporally smoothed by a factor of 4. Link: https://youtu.be/7LHpixvECHw

***Supplementary Video 3:*** Songbird organotypic slice exhibiting spontaneous, asynchronous calcium waves (30 DIV). Movie speed is increased by a factor of 4, and temporally smoothed by a factor of 4.Link: https://youtu.be/3qDAPGaBJa0

## ACKNOWLEDGEMENTS

The authors would like to thank the Gardner and Cruz-Martín lab, especially Ashley Comer for her insight and encouragement, Timothy Otchy for his useful comments, and Dawit Semu for providing excellent animal husbandry. Special thanks to D.S. Kim and L. Looger for providing the GCaMP6 DNA and to the GENIE project at Janelia Farm Research Campus, Howard Hughes Medical Institute. This work was supported by grants from NINDS **R24NS098536 and R01NS089679**.

